# Population history and gene divergence in Native Mexicans inferred from 76 human exomes

**DOI:** 10.1101/534818

**Authors:** María C. Ávila-Arcos, Kimberly F. McManus, Karla Sandoval, Juan Esteban Rodríguez-Rodríguez, Alicia R. Martin, Pierre Luisi, Viridiana Villa-Islas, Rosenda I. Peñaloza-Espinosa, Celeste Eng, Scott Huntsman, Esteban G. Burchard, Christopher R. Gignoux, Carlos D. Bustamante, Andrés Moreno-Estrada

## Abstract

Native American genetic variation remains underrepresented in most catalogs of human genome sequencing data. Previous genotyping efforts have revealed that Mexico’s indigenous population is highly differentiated and substructured, thus potentially harboring higher proportions of private genetic variants of functional and biomedical relevance. Here we have targeted the coding fraction of the genome and characterized its full site frequency spectrum by sequencing 76 exomes from five indigenous populations across Mexico. Using diffusion approximations, we modeled the demographic history of indigenous populations from Mexico with northern and southern ethnic groups splitting 7.2 kya and subsequently diverging locally 6.5 kya and 5.7 kya, respectively. Selection scans for positive selection revealed *BCL2L13* and *KBTBD8* genes as potential candidates for adaptive evolution in Rarámuris and Triquis, respectively. *BCL2L13* is highly expressed in skeletal muscle and could be related to physical endurance, a well-known phenotype of the northern Mexico Rarámuri. The *KBTBD8* gene has been associated with idiopathic short stature and we found it to be highly differentiated in Triqui, a southern indigenous group from Oaxaca whose height is extremely low compared to other native populations.

## Introduction

Comprehensive genome sequencing projects of human populations have demonstrated that a vast majority of human genetic variation has arisen in the past 10,000 years and is, therefore, specific to the continental and sub-continental regions in which they arose(Consortium 2012). As a result, the majority of rare variation contributing to disease burden is expected to be population specific and influenced by the local demographic history and evolutionary processes of each population(Gravel et al. 2011; Martin et al. 2017). Furthermore, it is recognized that there is a strong bias towards the inclusion of individuals of European descent in biomedical research, which is problematic for medical, scientific, and ethical reasons and should be counter balanced by including underrepresented populations in large genomic surveys of genetic variation(Bustamante, Burchard, and De la Vega 2011; Popejoy and Fullerton 2016).

Despite recent large-scale sequencing projects like the Exome Aggregation Consortium (ExAC)(Lek et al. 2016a) and gnomAD, which considerably expanded the knowledge on the patterns of protein-coding variation worldwide, little is known about the distribution of population-specific genetic variants that may underlie important evolutionary and biomedical traits of understudied groups. In particular, populations in the Americas of indigenous ancestry are expected to show exacerbated genetic divergence due to extreme isolation and serial founder effects during the continental peopling, leading to an increased fraction of population-specific variation (The 1000 Genomes Project Consortium 2015; Martin et al. 2017) that remains to be characterized. Present-day Mexico represents one of the largest reservoirs of Native American variation and a few studies have leveraged genotyping arrays and whole genome sequencing to characterize the genetic diversity and structure of the Mexican population. However, these studies used a limited number of markers or small sample sizes (Silva-Zolezzi et al. 2009; Moreno-Estrada et al. 2014; Romero-Hidalgo et al. 2017b), so there is a need to harness high coverage sequencing with population-level sampling. This will shed light on the consequences of functional variation in protein-coding genes as well as the adaptive and demographic processes that have shaped Native Mexican genomes

To fulfill this need, we sequenced the exomes of 78 individuals from five different indigenous groups from Northern (Rarámuri or Tarahumara, and Huichol), Central (Nahua), South (Triqui) and Southeast (Maya) Mexico. We characterized the protein-coding genetic variation from these populations to infer the broad demographic history of pre-Hispanic Mexico and to search for signatures of adaptive evolution.

## Results

### Genetic variation in 76 Native Mexican Exomes

According to previous genetic characterizations of indigenous Mexican groups, Mexico’s Native American ancestry is substructured into three major geographical components: Northern, Central and Southern(Gorostiza et al. 2012; Moreno-Estrada et al. 2014). In order to capture such substructure, we obtained protein-coding genetic variation from the sequences of 78 exomes from five Native Mexican (NM) populations representing all three major genetic regions of Mexico: Huichol (HUI, n=14), Maya (MYA, n=13), Nahua (NAH, n=17), Rarámuri (TAR, n=19) and Triqui (TRQ, n=15). Exomes were sequenced at an average depth of >90X (Supplementary figure 1). We used the Genome Analysis Tool (GATK)(McKenna et al. 2010a) to call variants jointly with an exome dataset including 103 Han Chinese (CHB) individuals from the 1000 genomes project (TGP)(The 1000 Genomes Project Consortium 2015). We jointly called with CHB exome data as we used the variants in this population for downstream analysis involving tests for selection in the NM groups. We identified 120,735 single nucleotide variants (SNV) and computed the genotype concordance between these and previously generated data from Affymetrix 6.0 (Moreno-Estrada 2014) and Axiom World IV (Galanter et al. 2014) SNP arrays available for the NM individuals. Concordance was above 93% for all individuals except for one TRQ and one HUI individual, which were excluded from all downstream analyses (Supplementary Fig 2) (Supplementary table S1). A predominance of Native American genetic ancestry in the remaining 76 NM individuals was corroborated with ADMIXTURE(Alexander, Novembre, and Lange 2009a) and principal components analysis (PCA) (Supplementary figure 3). Fifty-nine individuals displayed some non-Native ancestry ranging from 0.1% to 13%, and therefore we masked this fraction in the admixed exomes (see Material and Methods) for downstream analyses (Supplementary table S2).

After masking, a total of 58,968 SNV were retained in the 76 NM exomes with a transition/transversion ratio of 3.025. A subset of 4,181 SNVs was absent from public datasets (ExAC, TGP, and dbSNP v.142). The number of novel variant sites per exome ranged between 29 and 118 (median 84). Most of these novel SNVs are nonsynonymous (67.5%) and found at low frequencies: approximately 80% are singletons, while the rest are found at less than 5% frequency in the NM exomes (Supplementary Figs. 4-5). The number of singletons per population was 5,262 for the HUI (average per individual 405), 6,093 for the MYA (average per individual 469), and 8,108 for the NHA (average per individual 476), and 5,454 for the TAR (average per individual 287), and 5,166 for the TRQ (average per individual 369).

### Population history of Native Mexicans

We used the site frequency spectrum to infer the demographic history of four native Mexican populations: TAR, HUI, TRQ, and MYA. The NAH population was excluded from this analysis due to genetic substructure found within this linguistic group, which introduces noise in this type of analysis (see Discussion). We utilized a diffusion approximation approach implemented in the software δαδι (Gutenkunst et al. 2009) to infer the best-fit topology and demographic parameters of the four populations. The best fitting topology joins Northern populations together (HUI and TAR), as well as Southern populations (TRQ and MYA) stemming from a shared branch (Figure 1) (Supplementary table S3). The same topology is recovered when inferring split patterns with the program TreeMix (Supplementary figure S6) (Pickrell and Prichard 2012).

**Figure 1.**
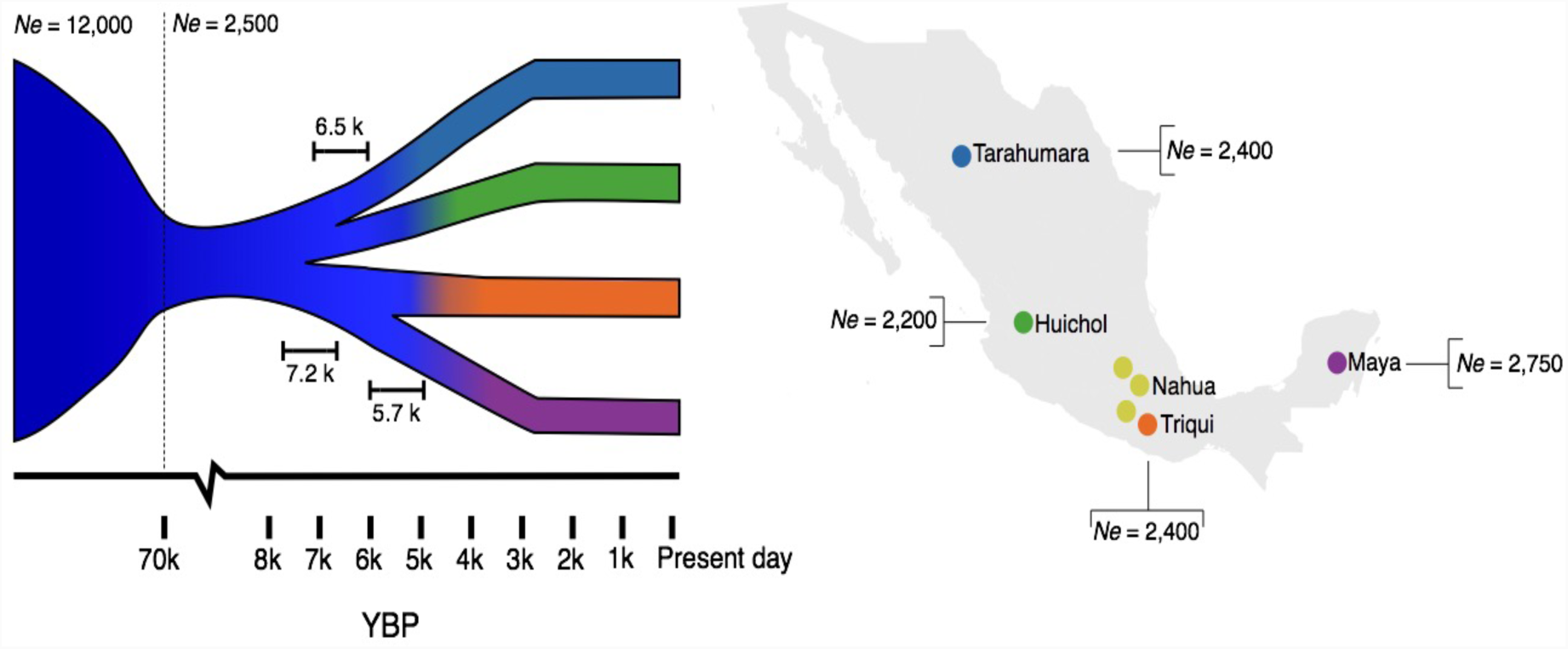
Sampling locations and inferred demographic model for NM populations. Inferred split times are shown on the demographic model and effective population sizes (*Ne*) are shown on the map. Each branch represents one of the populations used in the demographic inference; colors correspond to those shown in the map displaying the sampling locations of the participant NM. The Nahua were not included in the demographic inference (see discussion).

For all models, we fixed a population bottleneck around 70 kya, representing the Out of Africa bottleneck. Our best-fit model has an ancestral effective population (*Ne*) of 12,000 individuals for all NM, which is reduced to a *Ne* of 2,500 individuals after the bottleneck. We inferred that the split between northern (TAR and HUI) and southern (TRQ and MYA) Native Mexican populations occurred 7,200 years ago (Figure 1, Supplementary table S3). We find the two subsequent splits occurring within 1,500 years of each other: TAR and HUI diverged from each other 6,500 years ago, followed by the TRQ and MYA split 5,700 years ago (Figure 1, Supplementary table S3). We estimated all four populations to have similarly small *Ne*. The MYA has the largest Ne (2,750), followed by the TAR (2,400), TRQ (2,400), and HUI (2,200) (Figure 1, Supplementary table S3). 95% confidence intervals for these parameters were determined with 1000 bootstrapped replicates (Supplementary table S4). We note that this model assumes a constant population size since the last split. We were not able to estimate population growth rates due to the small sample size.

### Adaptive evolution in Native Mexicans

We used the called variants within the Native ancestry segments, to estimate an ancestry-specific population branch statistic (PBS)(Yi et al. 2010), which allowed us to control for non-indigenous admixture. This Fst-based statistic allows the identification of genes with strong differentiation between closely related populations since their divergence; it uses a third more distantly related population to detect changes affecting a specific population. We first estimated PBS by grouping all NM and using available exome data from the TGP to complete the topology with Han Chinese (CHB) and individuals of European ancestry (CEU). This allowed the detection of genes likely under selection in all Native Mexicans since divergence from the CHB.

We defined the genes in the 99.9^th^ percentile of the empirical distribution of the PBS values as being candidates of adaptive evolution (Figure 2 and Supplementary Table S5). Interestingly, some of these genes had previously been identified as targets of selection in other populations. These genes include *SLC24A5,* involved in skin pigmentation, *FADS3* involved in lipid metabolism, and *FAP*, which was previously suggested to be under adaptive archaic introgression in Peruvians (Racimo, Marnetto, and Huerta-Sánchez 2017) and Melanesians (Vernot et al. 2016). Of interest, three genes were involved in immune response: *SYT5* and *TBC1D10C*, involved in innate and adaptive immune response, respectively, and *MPZL1*, a surface receptor for a lymphocyte mitogen (R. Zhao and Zhao 2000; Runxiang Zhao et al. 2002). Several genes encode for cell membrane proteins (*GRASP, ADRBK1* and *SMPDL3A*) and cell-cell interactions (*MDGA2)*. The remaining genes were involved in spermatogenesis (*GMCL*) and angiogenesis (*NCKIPSD*).

**Figure 2.**
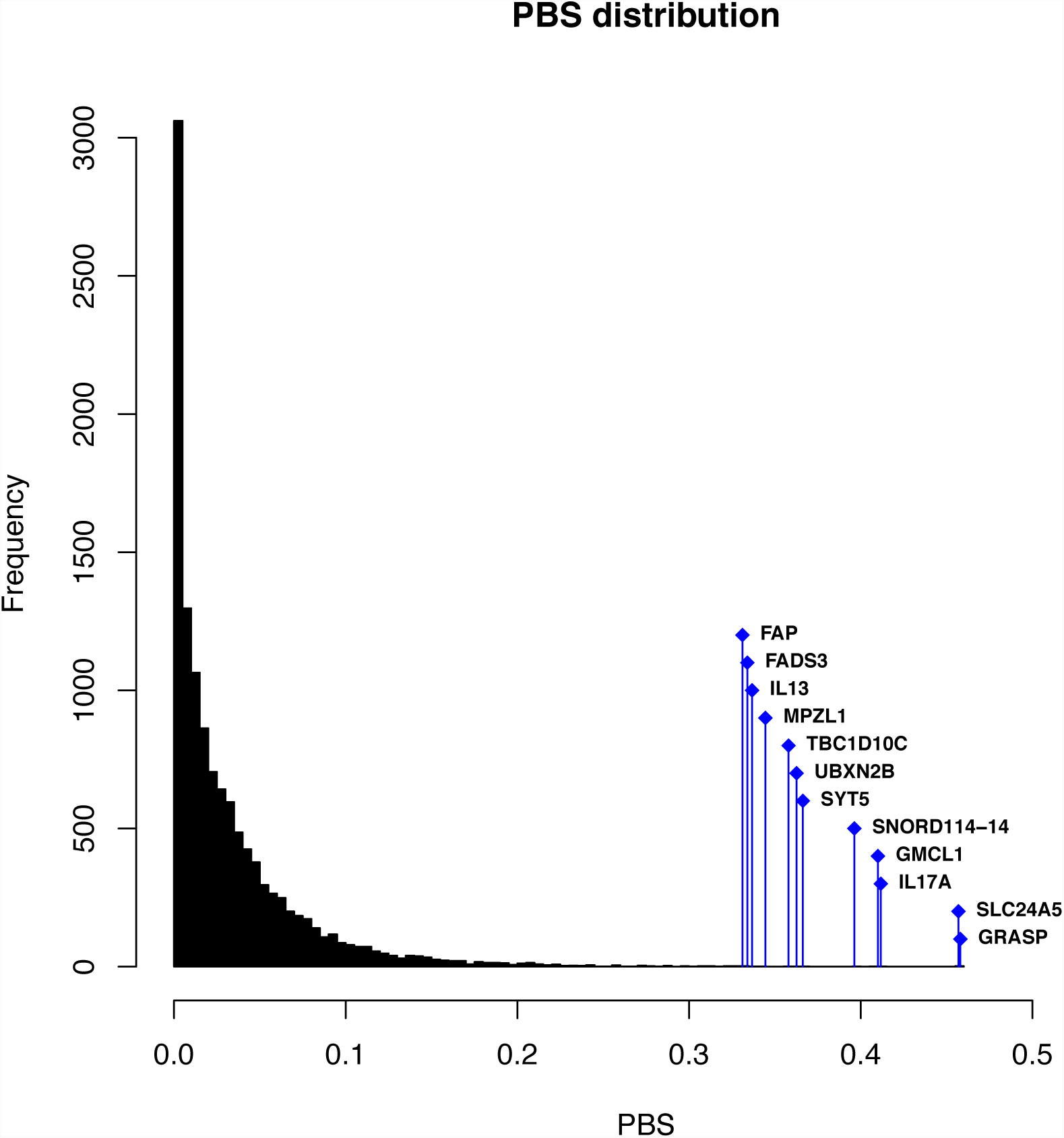
Distribution of gene-based population branch statistic (PBS) in NM. The x-axis shows the value of the PBS and the y-axis represents the frequency at which that value was observed among all NM. Genes displaying extreme (99.9^th^ percentile) PBS values are highlighted.

### Population-specific adaptive evolution within Mexico

To investigate genes under selection specific to each of the four NM populations for which we had a demographic model (HUI, MYA, TAR, and TRQ), we calculated PBS using the Han Chinese (CHB) as the third population in the form of NM1, NM2, CHB for all 12 combinations. To evaluate the significance of these PBS values we compared the observed data to PBS values obtained from simulations under the inferred demographic model (see methods). We were thus able to assign p-values to each gene and rank them by significance.

Genes with PBS values passing our significance threshold (p<10^−5^, Figure 3a-c) were identified only in HUI and TAR. In HUI, the genes with significant PBS were involved in innate immune response (*DUSP3*), cellular proliferation and differentiation (*KNC2*) and transcriptional repression of herpesvirus promoters (*ZNF426*). The genes in TAR (Figure 3c) included an open reading frame of unknown function (*C7orf25*) with high expression in testis, transformed fibroblasts and tibial nerve (GTex Version 7, Supplementary figure S7), and *FBX04,* involved in phosphorylation-dependent ubiquitination.

**Figure 3.**
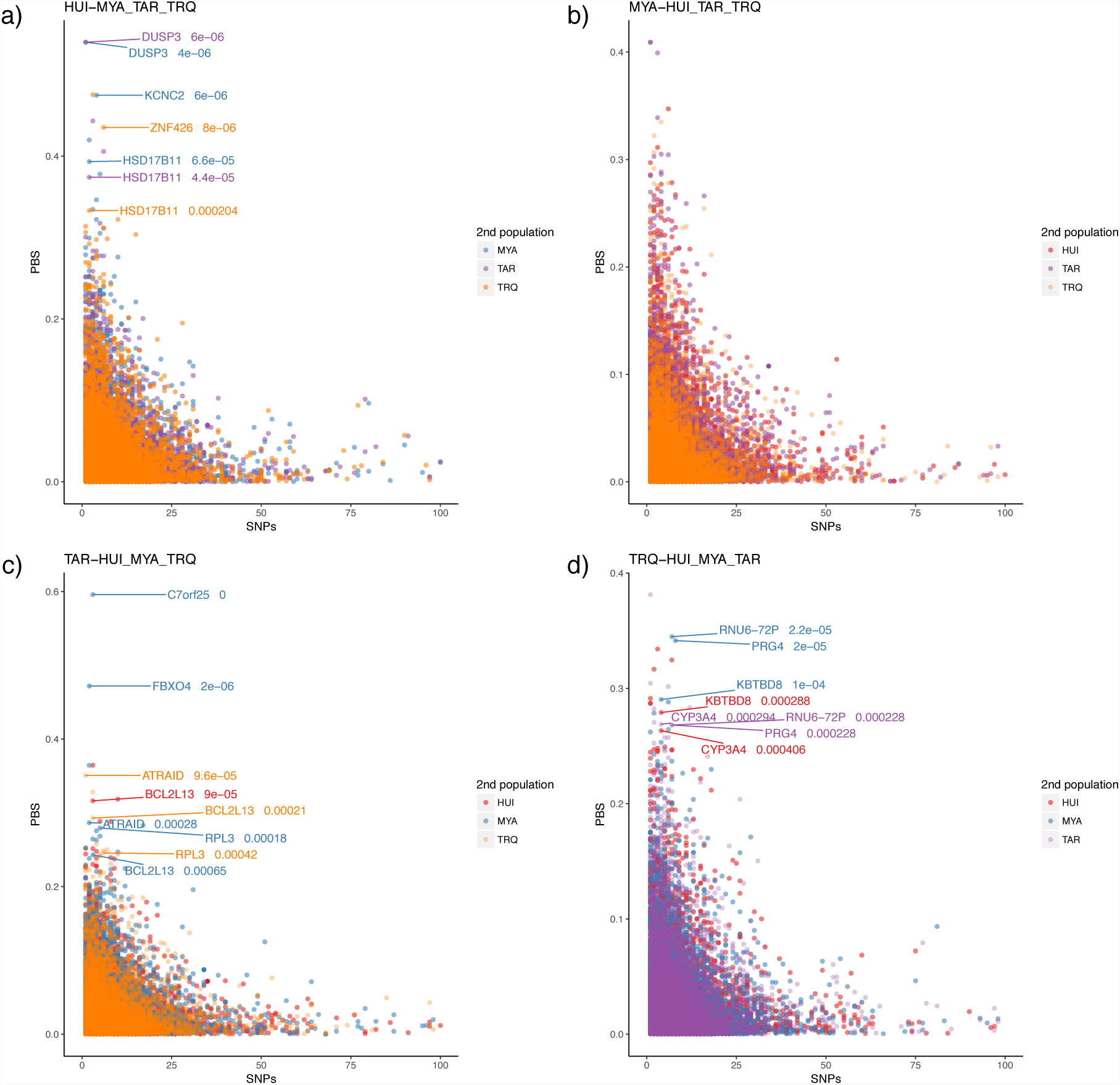
Population branch statistic (PBS) according to the number of SNPs in each gene for a) Huichol (HUI), b) Maya (MYA), c) Rarámuri (TAR), and d) Triqui (TRQ). Genes under likely adaptive evolution are shown with their corresponding p-value and. Colors represent the population used as the second population for the computation of the PBS (Han Chinese were always used as the third, distant population).

Furthermore, for each population we identified genes in the top 1% of the PBS distribution and in the lowest 1% of p values that were shared by at least two of the three possible pairwise comparisons between populations. This yielded nine additional candidate genes for adaptive evolution specific to different NM populations (Supplementary table S6). Of notice, two apoptosis-related genes were identified this way in TAR, *BCL2L13* (BCL2 like 13), and *ATRAID* (All-Trans Retinoic Acid-Induced Differentiation Factor). The former encodes for a pro-apoptotic protein that localizes in the mitochondria and is highly expressed in skeletal muscle (GTEx release V7) and is found in a locus previously associated to osteoarthritis (OA) risk in Mexican Americans (Coan et al. 2013). *ATRAID* (alternative name APR-3, apoptosis-related protein 3), is thought to be involved in apoptosis hematopoietic development and differentiation, and in positive regulation of bone mineralization and osteoblast differentiation (Zou et al. 2011).

Another noteworthy gene is *KBTBD8* (Kelch Repeat and BTB Domain Containing 8), which showed an extreme PBS in TRQ (Figure 3d). The encoded protein is involved in ubiquitination and is found in a locus previously associated to idiopathic short stature in Koreans (Kim et al. 2010). This is relevant because the TRQ (from the southern state of Oaxaca) display a particularly short stature and the SNV driving the selection signal (rs13096789) causes a nonsynonymous change classified as “possibly damaging” by PolyPhen. Lastly, we found gene *HSD17B11* (Hydroxysteroid 17-beta dehydrogenase 11) as an additional candidate of adaptive evolution in HUI. *HSD17B11*is a short-chain alcohol dehydrogenase that metabolizes secondary alcohols and ketones, and it has been suggested to participate in androgen metabolism during steroidogenesis.

To evaluate if genes showing extreme PBS values were enriched in any functional category or metabolic pathway we took the intersection of the genes with a p-value < 0.05 in all three pairwise comparisons for each NM population (see Methods). This way we compiled a list of 214 genes for HUI, 215 for MYA, 182 for TAR and 179 for TRQ (Supplementary tables S7-S10). We evaluated functional enrichment for the three Gene Ontology (GO) project categories: biological processes, cellular components, and molecular function (Ashburner et al. 2000; The Gene Ontology Consortium 2017), as well as pathway over-representation using the IMPaLa tool (Kamburov et al. 2011). We observed a functional enrichment with a significance of p < 0.05 in HUI involving lipid intestinal absorption (GO: 1904729, GO:0030300, GO: 1904478) (Supplementary table S11). Consistently, the IMPaLa pathway over-representation analysis revealed an enrichment of genes involved in lipid metabolism and transport, specifically the Statin pathway, for the same population (Pathway source: Wikipathways, p value (1.2e-05), Q value 0.0518).

## Discussion

We carried out the most comprehensive characterization of potentially adaptive functional variation in Indigenous peoples from the Americas to date. We identified in these populations over four thousand new variants, most of them singletons, with neutral, regulatory, as well as protein-truncating and missense annotations. The average number of singletons per individual was higher in Nahua (NHA) and Maya (MYA), which is expected given these two Indigenous groups embody the descendants of the largest civilizations in Mesoamerica, and that today Nahua and Maya languages are the most spoken Indigenous languages in Mexico (INEGI, 2015). Furthermore, the generated data also allowed us to propose a demographic model inferred from genomic data in Native Mexicans and to identify possible events of adaptive evolution in pre-Columbian Mexico.

### Demography

We propose, to our knowledge, the first demographic model that uses genetic data to estimate split times between ancestral populations within Mexico. By using the site frequency spectrum of neutral SNVs using a diffusion approximation approach, we inferred a split between northern and southern NM at approximately 6.7 to 8.2 KYA, followed by regional differentiation in the north at 5.9 to 7.5 KYA and 5.0 to 6.8 KYA in the south of Mexico (95% bootstrap CI, Supplementary table S4). This northern/southern split and a northwest to southeast cline is consistent with previous reports based on whole-genome and microarray genotype data from NM (Moreno-Estrada et al. 2014a; Romero-Hidalgo et al. 2017b). Furthermore, these split times are also coherent with previous estimates of ancestral Native Americans diverging ∼17.5-14.6 KYA into Southern Native Americans or “Ancestral A” (comprising Central and Southern Native Americans) and Northern Native Americans or “Ancestral B” (Raghavan et al. 2015; Rasmussen et al. 2014; Reich et al. 2012; Moreno-Mayar et al. 2018), and with an initial settlement of Mexico occurring at least 12,000 years ago, as suggested by the earliest skeletal remains dated to approximately this age found in Central Mexico (Gonzalez et al. 2003) and the Yucatán peninsula (Chatters et al. 2014). Studies on genome-wide data from ancient remains from Central and South America reveal genetic continuity between ancient and modern populations in some parts of the Americas over the last 8,500 years (Raghavan et al. 2015; Posth et al. 2018), though two ancient genomes from Belize (dated to 7,7400 BP and 9,300 BP, respectively) do not show specific allele sharing with present-day populations from that geographic area, instead they display similar affinities to different present-day populations from Central and South American populations, respectively (Posth et al. 2018). This suggests that by that time, the ancestral population of MYA was not yet genetically differentiated from others, so our estimates of northern/southern split at 7.2 KYA and MYA/TRQ divergence at 5.7 KYA fit with this scenario. Altogether these observations based on archaeological and paleogenomic data are consistent with our time estimates of population splits within Mexico, which involve a divergence of the Northern and Southern NM occurring at least two thousands years after the settlement and a divergence within these branches taking approximately between six hundred to two thousand years, respectively.

Regarding effective population sizes (*Ne*), we inferred an ancestral *Ne* of 2,500 for all NM, which is in line with a recent *Ne* estimate of 2,000 based on Markovian coalescent analyses of whole-genome data from twelve NM (Romero-Hidalgo et al. 2017b) as well as from Native ancestry segments in admixed Mexicans (Schiffels and Durbin 2014). Both studies show a low *Ne* around 2,000 sustained for the last 20 KY, in agreement with genomic and archaeological evidence pointing to a population bottleneck ca. 20 thousands years ago experienced by the Native American ancestors when crossing the Bering Strait into the Americas (Raghavan et al. 2015; Moreno-Estrada et al. 2014a; Goebel, Waters, and O’Rourke 2008).

In addition, our model inferred low *Ne* for present-day NMs ranging between 3,000 and 3,800. The NM with the largest *Ne* was the MYA (95% CI: 2,400-3,400) followed by TAR (95% CI: 2,200-2,800), HUI (95% CI: 1,900-2,600), and TRQ (95% CI: 2,100-2,900) (Supplementary table S4). Using runs of homozygosity Moreno-Estrada et al. (2014) inferred slightly higher variation in *Ne* among different indigenous groups, but with overlapping confidence intervals. On the other hand, using whole-genome data Raghavan et al. (2015) inferred a HUI *Ne* to a similar 2,500. Of notice, the census size of these populations is also the largest for the MYA, and TAR (INEGI 2015). These observations are noteworthy since low *Ne* combined with founder effects can exacerbate the disproportionate accumulation of deleterious and clinically relevant variants in the population(Belbin et al. 2018), and indeed these two populations also displayed the largest numbers of singleton and nonsynonymous SNVs.

One caveat of our demographic inference is that we failed to include the NHA in the model. Initial tests including this population resulted in extremely high Ne and ambiguous location in the tree. Furthermore, ADMIXTURE analyses showed NHA are constituted by components from multiple populations within each individual (Supplementary figure S3a), an observation made also in a recent study by Romero-Hidalgo et al. (2017). This likely reflects NHA being genetically heterogeneous as a consequence of their past history involving continuous colonization and extended domination of multiple distinct groups by the Nahua-speaking Aztec empire right before European colonization (Brumfiel 1983; Romero-Hidalgo et al. 2017b).

Overall, our inferences describe, to our knowledge, the most detailed demographic history model based on genetic data for NM to date. We caution, however, that as with any inferred demographic model, the assumptions have certain caveats that could lead to errors. Specifically, our model assumes constant population sizes since the most recent split. Certainly the availability of genome-wide data from present day, as well as from ancient populations from throughout Mexico spanning these time frames, will contribute to draw a more refined picture of past population history and genetic structure.

### Targets of adaptive evolution in NM

We implemented an ancestry-specific approach of the widely used FST-based Population Branch Statistic (PBS) to identify genes with strong differentiation in all NM since their divergence from CHB, as well as genes differentiated in each studied population. The first approach revealed selection signals previously found in Native Americans and other populations, as well as genes not previously identified to be under selection.

One remarkable instance is *FADS3*, a gene in the FADS (fatty acid desaturases) genes cluster in chromosome 11, which also includes *FADS1* and *FADS2*. These genes are involved in the metabolism of omega-3 polyunsaturated fatty acids, and have been found to harbor strong signals of positive selection in Arctic populations (Fumagalli et al. 2015) and throughout the Americas (Amorim et al. 2017). It has been proposed that this signal derives from a strong selective pressure on the ancestors present-day Native Americans (Amorim et al. 2017) who had to adapt to extreme cold weather and food availability during the Beringia standstill (Tamm et al. 2007).

We also identified a strong differentiation on *FAP* (Fibroblast Activator Protein Alpha). The locus harboring this gene, together with *IFIH1*(interferon induced with helicase C domain 1) is suggested to be under adaptive archaic introgression in Peruvians from the TGP (PEL), a population with a high proportion of Native American genetic ancestry, (Racimo, Marnetto, and Huerta-Sánchez 2017), and Melanesians (Vernot et al. 2016). This locus has been associated with type 1 diabetes (Liu et al. 2009) and susceptibility to diverse viral infections(Fumagalli et al. 2015). The fact that this locus also has an adaptive signal in NM is consistent with a previous study that suggested that the loci harboring *IFIH1* suffered recent positive selection in South Americans (Fumagalli et al. 2010). We confirmed that this haplotype is present in NM by comparing the Neandertal haplotype with whole genome sequence data (Marnetto and Huerta-Sánchez 2017) available for other Native Americans from an independent study (Romero-Hidalgo et al. 2017b). The archaic haplotype was found in 7 out of 24 chromosomes in TAR and MYA individuals as well as in other NM (Tepehuano, Totonaco and Zapotec) (Supplementary Figure S8).

*SLC24A5* (solute Carrier family 24 Member 5) has also been previously identified as target of selection. This gene is involved in melanogenesis and has been vastly studied in European populations, where it displays one of the strongest signals of selection in humans. Specifically, the derived allele of SNP rs1426654 within this gene leads to a decrease in skin pigmentation. However, the ancestral allele is nearly fixed in NM populations, driving the extreme PBS signal and suggesting either a relaxation of selection or an adaptive event favoring the ancestral state (the derived allele is fixed in CEU, and found at 0.03 and 0.007 frequencies in CHB and NM, respectively).

In addition, we found several genes with extreme PBS values involved in immunity and defense against pathogens (*SYT5, TBC1D10C* and *MPZL1*). Selection on these genes could be explained by the strong selective pressure posed by pathogens brought by Europeans on the Native population during colonization; it is estimated that up to 90% of the Native population died as a consequence of infections during this period (Acuna-Soto et al. 2002). Lastly, three genes in this set were cell-surface and signal transduction genes (*GRASP, ADRBK1* and *SMPDL3A, MDGA2*).

Regarding population-specific signals of adaptive evolution, we identified few genes with significant (p<10^−5^) PBS values in TAR and HUI only. These genes were also involved in innate immune response (*DUSP3*) and repression of herpesvirus transcription (*ZNF426*), cellular proliferation and differentiation (*KNC2*), and ubiquitination (*FBX04)*. When we expanded our search to consider genes above the significance threshold, but with consistent extreme PBS (top 1%) in two or more population comparisons, we identified some interesting instances in TRQ, HIU and TAR, which we speculate could be related to some characteristic traits in these populations.

Anthropometric studies have revealed that the TRQ (together with other Indigenous groups in Oaxaca, and neighboring states of Veracruz and Chiapas) exhibit the lowest average stature values in Mexico (females mean 142.5cm, males mean 155.1)(Faulhaber 1970). This observation becomes relevant as one of the genes (*KBTBD8*) identified as likely being under adaptive evolution in this population, lies within a locus previously associated to idiopathic short stature in Koreans (Kim et al. 2010). Regarding HUI, we found a gene (*HSD17B11*) involved in the metabolism of steroids and retinoids, with high expression in tissues related to steroidogenesis (adrenal gland and testis) and detoxification (liver, lung, kidney and small intestine)(Lundová et al. 2016). Interestingly, in the region of Mexico where HUI are from, the ceremonial intake of peyote cactus (*Lophophora williamsii)* is a cultural tradition that traces back to centuries. The psychoactive compound in peyote, the alkaloid mescaline, is metabolized by the liver enzymes and can cause severe toxicity when consumed in high amounts. Furthermore, the pathway enrichment analysis in this population returned genes APOA1, APOA2, APOA4, APOA5, ABCG5 (Supplementary Table S12), involved in regulation of intestinal cholesterol absorption. Variants in some of these genes have been associated to high levels of LDL and total cholesterol (Bandarian et al. 2013; the ENGAGE Consortium et al. 2009)(Bandarian et al., 2013; the ENGAGE Consortium et al., 2009)(Bandarian et al. 2013; the ENGAGE Consortium et al. 2009)(Bandarian et al. 2013; the ENGAGE Consortium et al. 2009) as well as high triglyceride levels (Pennacchio et al. 2002; Zhu et al. 2014; Ouatou et al. 2014), both factors leading to cardiovascular disease. The derived allele of rs3135506 in *APOA-5* has the highest frequency in HUI compared to the other populations from the study and the TGP1000 (Supplementary Table S12). This missense mutation (S19W) has been associated with increased triglyceride levels and elevated risk of developing coronary artery disease in several populations including one labeled as “Hispanic” (Pennacchio et al. 2002; Zhu et al. 2014; Ouatou et al. 2014). Moreover, HUI has the highest frequency of the missense mutation rs6756629 in *ABCG5*, which has been associated to increased total cholesterol and LDL (the ENGAGE Consortium et al. 2009), a risk factor for coronary heart disease. Together, these observations point some possible adaptation to a low cholesterol-lipid diet (such as a reduced meat consumption) or a manner to regulate the intake of lipids from animal source foods.

Lastly, two genes (*BCL2L13* and *ATRAID*), in TAR have annotations related to joint and bone physiology, namely osteoarthritis, bone mineralization and osteoblast differentiation, which could be related to the outstanding physical endurance in the Rarámuri. Interestingly, a recent study detected an enrichment of genes harboring novel promoter and missense variants with pathway and gene ontology (GO) annotations related to musculoskeletal function (Romero-Hidalgo et al. 2017b) in the same population. In agreement with this, we recapitulated similar observations when looking for GO enrichment in novel nonsynonymous variants in our 19 TAR exomes (Supplementary table S13). These two approaches represent independent evidence from both highly diverged and novel functional variation that converge in musculoskeletal traits as the potential underlying mechanism for Raramuri’s endurance. Taken together, these observations could imply that this cultural trait has imposed a selective pressure on this population. However, additional in-depth studies in the Raramuri incorporating genomic and detailed phenotype data are needed to disentangle the genetic architecture and the molecular pathways behind this complex trait.

In conclusion, we generated a rich catalogue of Native American genetic variation from Mexican populations, the analysis of which has yielded novel estimates for ancestral population splits as well as candidate genes likely under adaptive evolution in both the general NM population and in specific NM groups. Our demographic inference is consistent with previous archaeological and genetic knowledge on the peopling of the Americas, while adding temporal resolution to the population dynamics occurring thousands of years ago in the Mexican mainland. This demographic model also allowed us to compare the estimated PBS values of genes against a simulated null distribution under such model, and to identify the instances with significant extreme values. Genes with extreme values in specific populations have annotations that hint a likely role in characteristic phenotype or cultural practices in the NM included in this study. However, it remains to be tested, if these high values indeed derive from adaptive events and if these adaptations are in fact involved with the observed traits in these NM populations.

## Materials and methods

### Samples

Most of the samples sequenced in this study were previously collected and sampling procedures are described in (Moreno-Estrada et al. 2014a). Specifically, a subset of samples from 4 of the studied populations were selected for having the highest proportions of Native American ancestry according to Affymetrix 6.0 SNP array data generated therein. After filtering for DNA quality control a total of 19 Tarahumara (Chihuahua), 13 Huichol (Jalisco), 15 Triqui (Oaxaca), and 12 Maya (Quintana Roo) individuals were included in this study. Additionally, 18 Nahua samples from three sampling locations in Central Mexico previously genotyped with Affymetrix Axiom World Array IV (Galanter et al. 2014)were selected for exome sequencing based on their proportions of Native American ancestry and passing DNA quality control. In both sampling schemes, Institution Review Board (IRB) approval was obtained from Stanford University, and individuals were consented according to the approved protocol. All individuals gave written consent. DNA was extracted from blood and ethnographic information including family, ancestry, and place of birth were collected for all individuals.

### Exome sequencing

Exome regions were captured using the Agilent SureSelect 44Mb human all-exon array v2 for the 76 individuals. Genotype data from previous studies (Moreno-Estrada et al. 2014b; Galanter et al. 2014a) was available for these individuals (Affymetrix 6.0 SNP array data for Huichol, Maya, Tarahumara, and Triqui, and Axiom World Array IV data for the Nahua).

Each individual was sequenced in a 5-plex library on an Illumina HiSeq 2000 producing 101-bp paired end reads. Reads were processed according to a standard pipeline informed by the best-practices described by the 1000 Genomes Project. Briefly, reads were mapped to the human reference genome (hg19) using bwa (version 0.6.2). Duplicate read pairs were identified using Picard (http://picard.sourceforge.net). Base qualities were empirically recalibrated and indel realignment was performed jointly across all samples using the Genome Analysis Tool Kit (GATK, version 1.6) (McKenna et al. 2010b). Variants were filtered to the exome capture region.

### Masking of non-Native genomic segments

To corroborate the Native American ancestry of the individuals we combined the data with variants from the 1000 Genomes Project (1000G) to perform principal components analysis.

Additionally, we estimate individual ancestries from the genotype data with the maximum-likelihood–based clustering algorithm ADMIXTURE (Alexander, Novembre, and Lange 2009b) to identify those samples with detectable contribution of European or African genetic ancestry. This fraction was masked to avoid their inclusion in the downstream analysis. To mask non-native regions in the exome data, the available genotype data for the individuals in the study was merged with a reference panel and processed as follows. We used the genotype data available for HUI, MYA, TAR and TRQ (Moreno-Estrada 2014) merged it with genotype data from the same array from 30 individuals with a 100% indigenous ancestry, as well as with 30 CEU, and 30 YRI from the HapMap Project to serve as reference panel. For NHA we combined the genotype data available for the admixed individuals(Galanter et al. 2014c) with genotype data of 20 Native American individuals genotyped with the same array, namely Affymetrix Axiom World Array IV (also known as LAT array for its informativeness in Latino populations), as well as with 20 CEU and 20 YRI, to serve as a reference panel..

Genotype data for each set was then phased using SHAPEIT (Version 2) (O’Connell et al. 2014) with default parameters. Local ancestry was estimated for the resulting haplotypes using the ‘PopPhased’ routine of RFMix(Maples et al. 2013) with parameters –correct-phase (for phase correction) and –G15 (to assume 15 generations since admixture). Local ancestry calls were then used to mask (make missing) the sites in the exome vcf files that were not part of homozygous Native American-ancestry blocks.

### Variant analysis and annotation

The VCF file was annotated using the tool ANNOVAR (version 2015Jun17)(K. Wang, Li, and Hakonarson 2010) with the following reference datasets: refGene, esp6500siv2_all, 1000g2015aug_all, exac03, avsnp142 and ljb26_all (see supplementary table XX for description of each dataset).

The script table_annovar.pl was used with the following parameters:

table_annovar.pl –vcfinput $pop.vcf annovar_humandb/ --buildver hg19 -- out $pop –remove --protocol refGene,esp6500siv2_all,1000g2015aug_all,exac03,snp142,clinvar_20160302,d bscsnv11,dbnsfp30a --operation g,f,f,f,f,f,f,f --nastring.

New variants were defined as those not present in ExAC (Lek et al. 2016b), NHLBI Exome Sequencing Project (ESP) (https://esp.gs.washington.edu), 1000g (The 1000 Genomes Project Consortium 2015b), and dbSNP142 datasets. These were discovered in the entire dataset of 76 exomes and also per population. The annotation of these new variants was retrieved and classified according to their potential effect on transcripts.

### Demographic inference

For demographic inference we used the exome sequencing data from the four NM populaitons HUI, MYA, TAR and TRQ, as well as data from Han Chinese (CHB) from TGP(The 1000 Genomes Project Consortium 2015b). Non-native ancestry was masked in each native Mexican sample. To include only neutral sites in the analysis, we limited to 4-fold and intronic sites determined via SNPEff(Cingolani et al. 2012). Inference was made on the unfolded site frequency spectrum (SFS). We used the panTro4 reference sequence as an outgroup and implemented a context-dependent correction for ancestral misidentification(Hernandez, Williamson, and Bustamante 2007). We estimate the chimpanzee reference genome to have a 0.012 divergence from the human reference (hg19) in our target regions. After removing triallelic sites and sites with a missing outgroup allele, we produced a callable sequence length of 8,889,201bp.

#### Dadi

The demographic model was inferred via an approximation to the forward diffusion equation implemented in δαδι(Gutenkunst et al. 2009). This approach infers the best-fitting parameters given a specific demographic model and calculates the log-likelihood of the model fit based on a comparison of the expected to observed site frequency spectrum. δαδι can handle a maximum of three populations and has difficulty optimizing with more than two populations. Due to this, we optimized over the pairwise population demographies (extending the approach from(Moreno-Estrada et al. 2013)). We projected population allele frequencies to the following numbers of chromosomes: TAR 26; MYA 14; TRQ 16; HUI, 16.

We fixed a population bottleneck in the ancestral population 70 kya (δαδι parameter: t=0.09). This parameter was fixed as it has been estimated in previous studies(Li and Durbin 2011) and because δαδι often has difficulties inferring the time and size of a bottleneck. As the expected SFS of multiple bottlenecks looks nearly identical to the SFS of one bottleneck (with a different magnitude), we expect this bottleneck to encompass the loss of diversity in the out-of-Africa expansion and the crossing of the Bering strait (similar to Raghavan et al. 2015). We utilized the topology inferred via Treemix (Supplementary Figure S6 a, Pickrell and Prichard 2012), and confirmed this as the best-fitting topology in δαδι (Supplementary table S4). To convert best-fit parameters to interpretable values, we assumed a generation time of 29 years and a mutation rate of 1.25e-8 mutations per base pair per generation. Then we set a topology and optimized over the best-fit for all pairwise population split models.

Confidence intervals were determined via 1000 bootstrapped replicates. To make these replicates, we divided the genome into 500 kb blocks and removed the blocks that contained no target regions. Then we randomly sampled blocks with replacement and inferred δαδι parameters.

### Positive Branch Statistic

We calculated the Positive Branch Statistic (PBS) as in (Yi et al. 2010)allele frequencies of the derived alleles per population and then calculated pairwise FST values for each gene using Reynold’s FST formula (Reynolds, Weir, and Cockerham 1983). Only sites with at least 10 chromosomes per population were included in the calculation. Also, only sites that were polymorphic in at least one of the three populations were considered. PBS values were then computed for each gene using the formula:

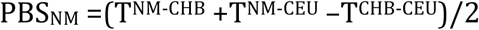

when considering all NM as a single population, and:

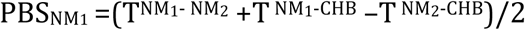

for each NM population (yielding twelve possible comparisons)

Where:

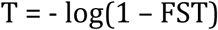

To evaluate the significance of the PBS values, we compared them to a null distribution of simulated neutral sequences and followed the method of Yi et al. (2010). We simulated sequences in *ms*(Hudson 2002) with our inferred 4-population NM demography and used Han Chinese (1000 Genomes, CHB) as an outgroup. We estimated the Chinese-Mexican split time by averaging the inferred split time and bottleneck *Ne* between the CHB and each native Mexican population.

Simulated allele frequencies where generated for 700k genes with 1 to 80 SNPs per gene. PBS values on the simulated data were then calculated using the same filters and formula as for the observed data. Because of filters (at least 10 chromosomes per population and only polymorphic sites), some of the simulated sites were disregarded causing a change in the number of simulations for each SNPs/gene bin. Because of this we randomly subsampled 500k simulations for each SNPs/gene bin category, which covered all bins between 1 to 71 SNPs/gene.

Observed PBS values were then compared to the simulated values. A p-value was calculated by observing the fraction of simulated PBS values larger than the observed PBS. For example, if there was only one simulated PBS value larger than the observed one, the p-value corresponded to a p-value of 0.000002 (1/500,000). For genes with 1 to 71 SNPs/gene observed values were compared to their respective simulated bins. Observed PBS values for genes with >71 SNPs, were compared to the simulated PBS distribution of 71 SNPs/gene.

### Functional enrichment analysis

To perform the enrichment analysis we first defined an intersecting subset of genes for each population considering only the genes in all pairwise comparison showing extreme PBS values (p-value < 0.05) (e.g. the intersection of the subsets TAR-HUI, TAR-MYA and TAR-TRQ generates list of intersecting genes for the Tarahumara.

For GO enrichment we used the online tool in http://www.geneontology.org/page/go-enrichment-analysis. We analyzed each of the four gene lists with the three GO categories (biological processes, cellular components and molecular function) using Bonferroni correction. (ran on 28/06/17)

To test for enrichment in GO categories among genes with novel missense SNV we used WebGestalt online tool(J. Wang et al. 2013) as reported in (Romero-Hidalgo et al. 2017a).

We ran a pathway over-representation analysis on the same gene sets using the IMPaLA online tool(Kamburov et al. 2011) available at: http://impala.molgen.mpg.de (ran on Sept/2017) and considered only pathways with a Q value less than 1. A score of 1 implies 100% of the results with that corresponding p-value are false positives, in this manner we took into account lower values.

## Acknowledgements

We thank the participants and volunteers who donated DNA samples and the fieldwork teams led by Hector Rangel, Victor Acuña, and Leonor Buentello on the various sampling expeditions. We thank M.C. Yee and M. L. Carpenter for technical support and E. Huerta-Sánchez and Diego Ortega del Vecchyo for input on early versions of the manuscript. We deeply thank the generous support from the Beijing Genomics Institute (BGI) for contributing with sequencing capacity, and the Stanford Center for Computational, Evolutionary and Human Genomics (CEHG) for supporting the initial stages of this project. We thank IT support from Luis Aguilar, Alejandro de León, Carlos S. Flores, and Jair García of the Laboratorio Nacional de Visualización Científica Avanzada at UNAM. This work has been supported by Mexico’s CONACYT Basic Research Program (grant number CB-2015-01-251380 awarded to A.M.-E.) and the International Center for Genetic Engineering and Biotechnology (ICGEB) Grant CRP/MEX15-04_EC awarded to A.M.-E. M.A.-A. was partially supported with a fellowship from the 2012 George Rosenkranz Prize for Health Care Research in Developing Countries awarded to A.M.-E. MCAA’s laboratory is supported by Programa de Apoyo a Proyectos de Investigación e Innovación Tecnológica - Universidad Nacional Autónoma de México grant IA206817 and CONACYT Infrastructure grant 26944.

