## Supplementary Figures and Tables for "Population history and gene divergence in Native Mexicans inferred from 76 human exomes"

### Supplementary figures index

**Supplementary figure S1** – Average exome depth-of-coverage per NM individual
**Supplementary figure S2** – Concordance between exome SNVs and genotyping array data per NM individual
**Supplementary figure S3** – ADMIXUTRE and Principal Components Analysis (PCA) of NM exome data
**Supplementary figure S4** – Distribution of allele frequencies of known and novel SNVs in NM exome data.
**Supplementary figure S5** – Functional and frequency distribution of known and novel SNVs in NM.
**Supplementary figure S6** – Tree topology inferred by TreeMix
**Supplementary figure S7** – GTEX plot of C7orf25
**Supplementary figure S8** - FAP/IFIH1 introgressed locus haplotypes.

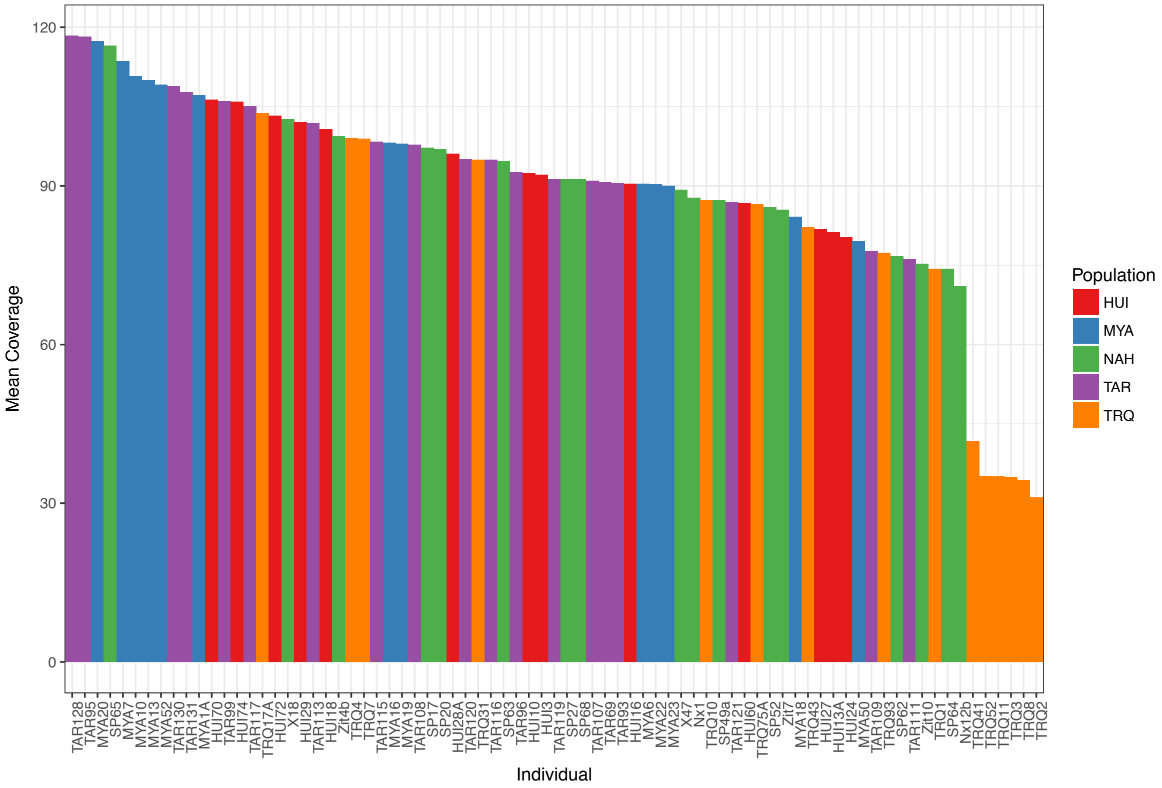


**Figure S1.**
Average exome depth-of-coverage per NM individual. Individual IDs are shown in the x-axis, and the color represents the NM population they belong to. Huichol (HUI), Maya (MYA), Nahua (NHA), Rarámuri (TAR), and Triqui (TRQ).

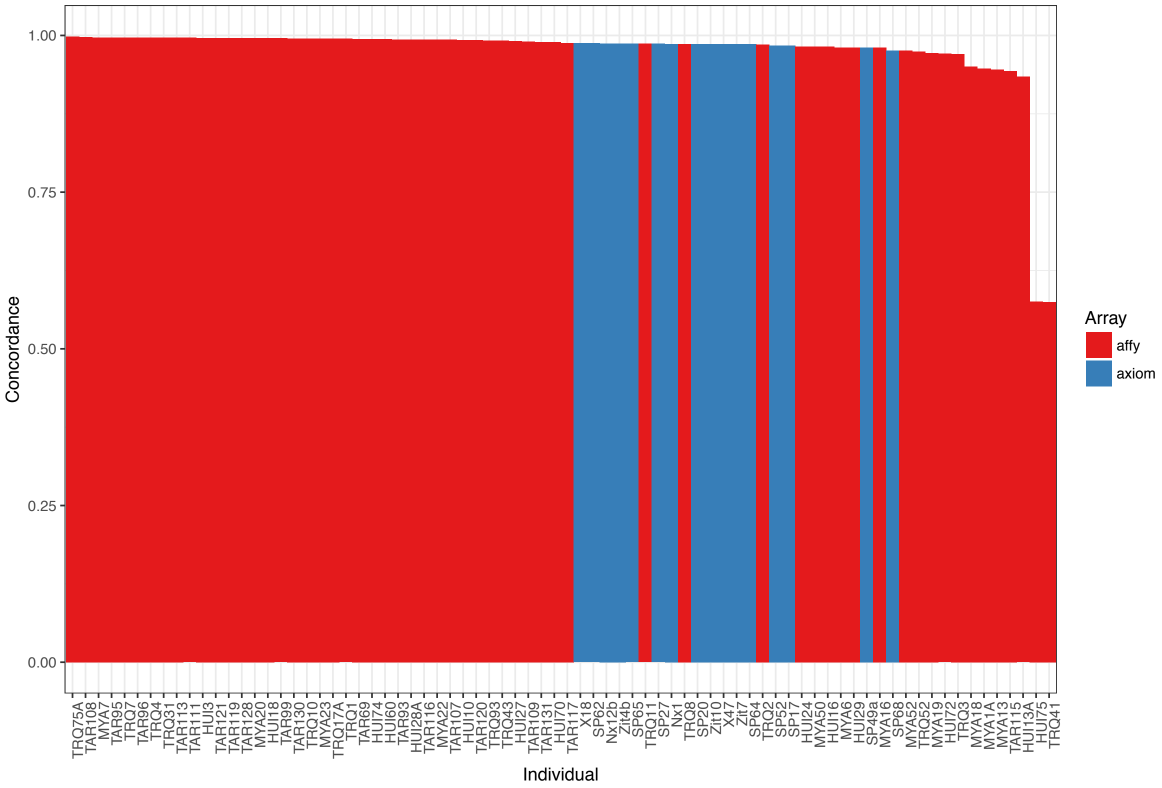


**Figure S2.**
Concordance between exome SNVs and genotyping array data per NM individual. Individual IDs are shown in the x-axis, and the color represents type of microarray used to generate the genotype data. Individuals genotyped with the Affymetrix 6.0 array are shown in red (Moreno-Estrada, 2014) while in individuals genotyped Affymetrix Axiom World Array IV are shown in blue (Galanter et al. 2014).

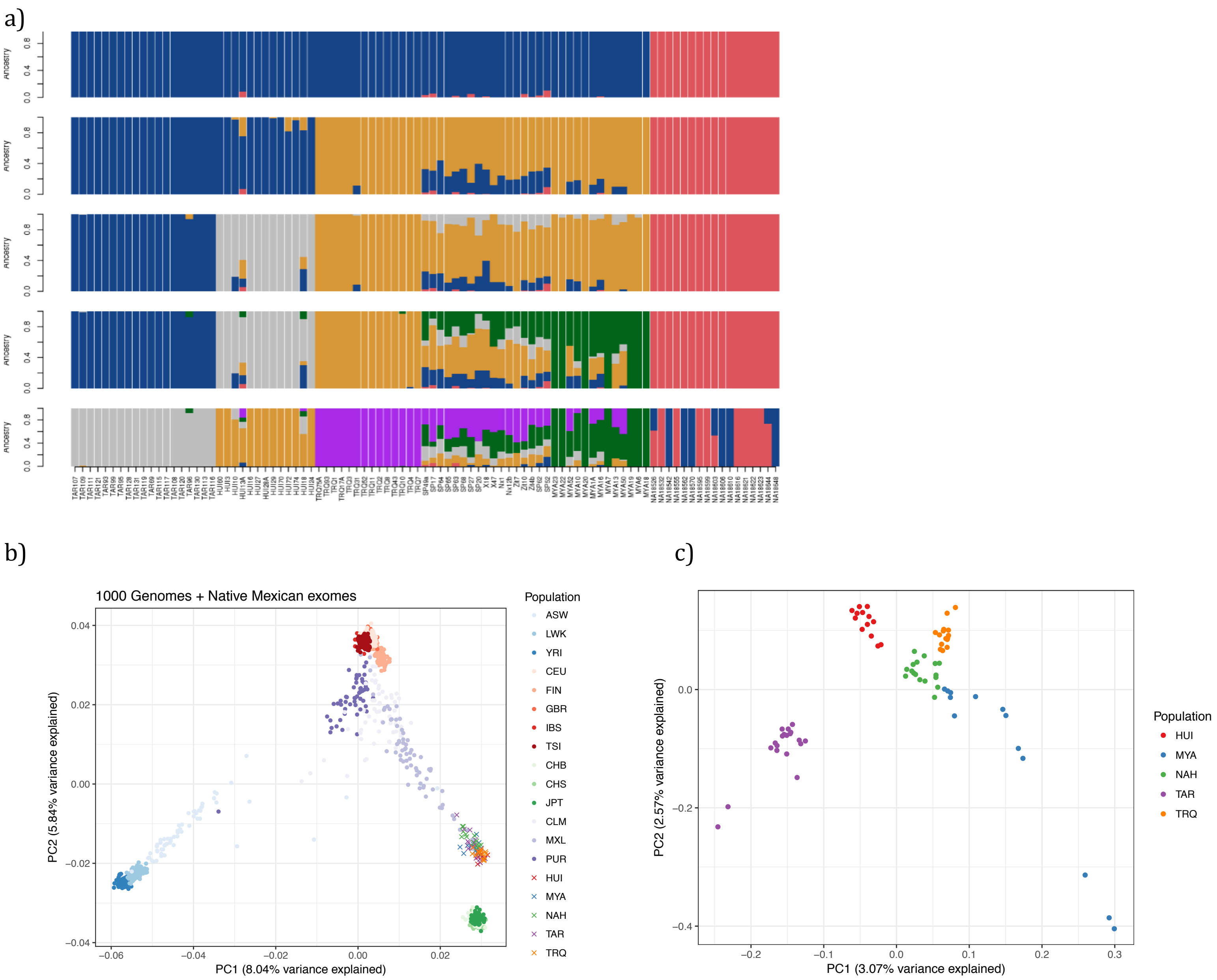
a)
Ancestry
0.0
0.4
0.8
b)
1000 Genomes + Native Mexican exomes
PC1 (8.04% variance explained)
PC2 (5.84% variance explained)
Population
ASW
LWK
YRI
CEU
FIN
GBR
IBS
TSI
CHB
CHS
JPT
CLM
MXL
PUR
HUI
MYA
NAH
TAR
TRQ
c)
PC1 (3.07% variance explained)
PC2 (2.57% variance explained)
Population
HUI
MYA
NAH
TAR
TRQ

**Figure S3.**
ADMIXUTRE and Principal Components Analysis (PCA) of NM exome data. **a)** ADMIXTURE plot with K=2 to K=6 (from top to bottom) for all NM. **b)** PCA of NM and worldwide populations from the 1000 genomes project (TGP) shows a clustering of the NM in at the Native American (NA) extreme of the Europe-NA cline displayed by the Mexicans (MXL) from the TGP, and **c)** PCA of NM exome-only data shows population substructure within NM, which separates northern Rarámuri (TAR) and Huichol (HUI), central Nahua (NHA), and southern groups Triqui (TRQ) and Maya (MYA).

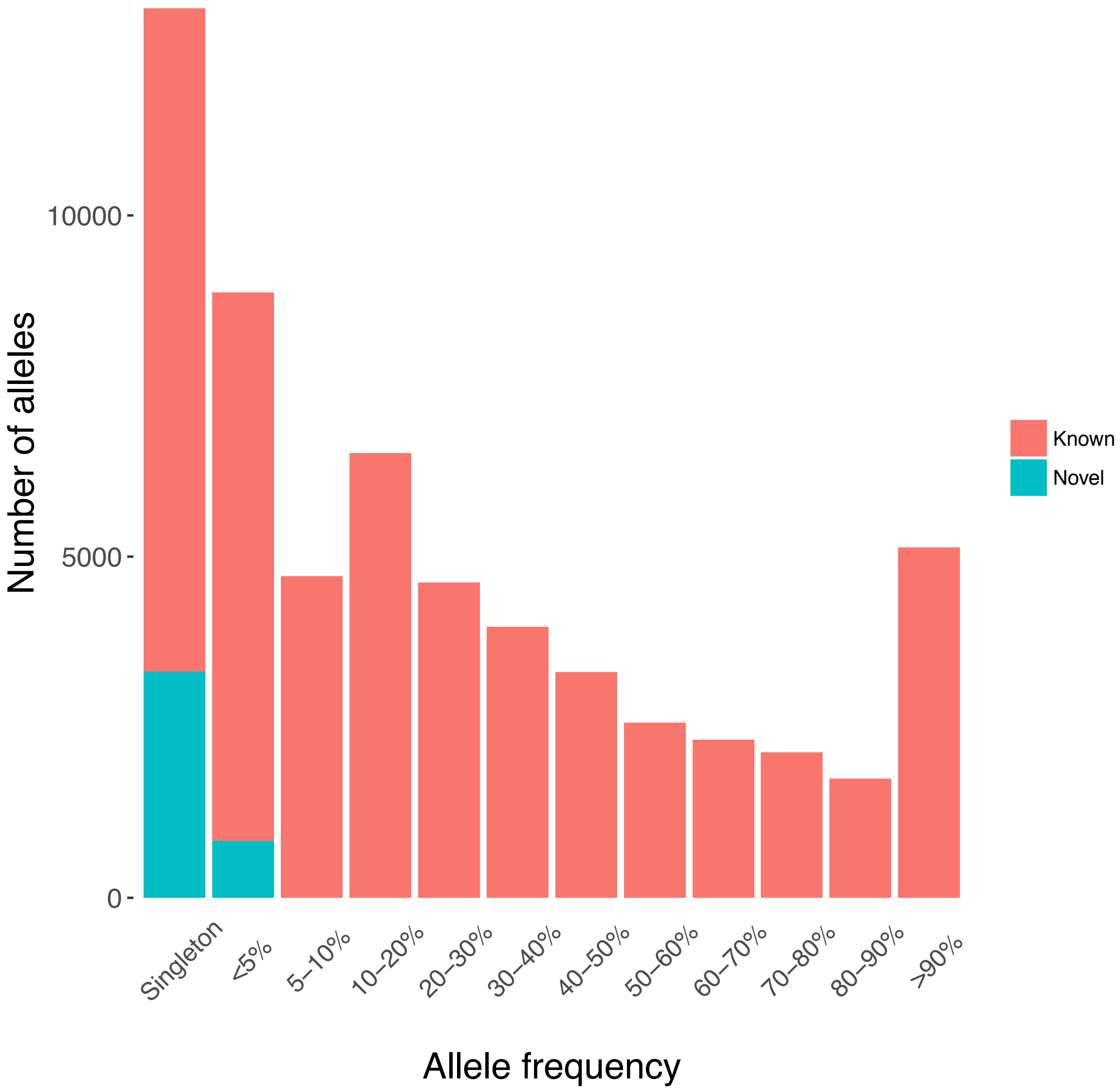


**Figure S4.**
Distribution of allele frequencies of known (pink) and novel (turquoise) SNVs in NM exome data. Most novel alleles are found as singletons or at very low (<5%) frequency.

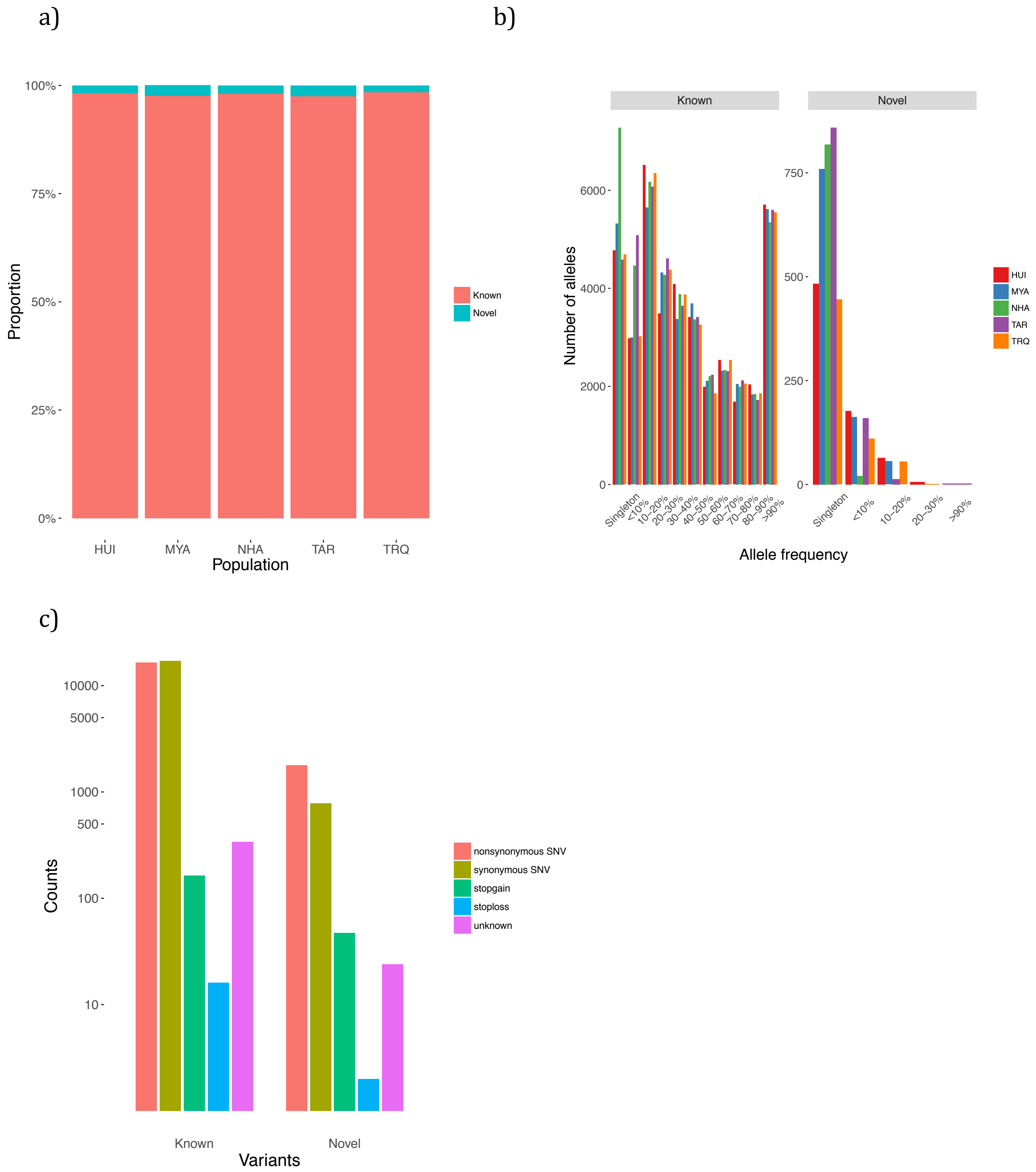


**Figure S5.**
Frequency distribution and functional annotation of known and novel SNVs in NM. **a)** Breakdown of novel and known SNVs by NM population. **b)** Distribution of allele frequencies of known known and novel SNV's per population. **c)** Functional annotation of novel and known SNVs.

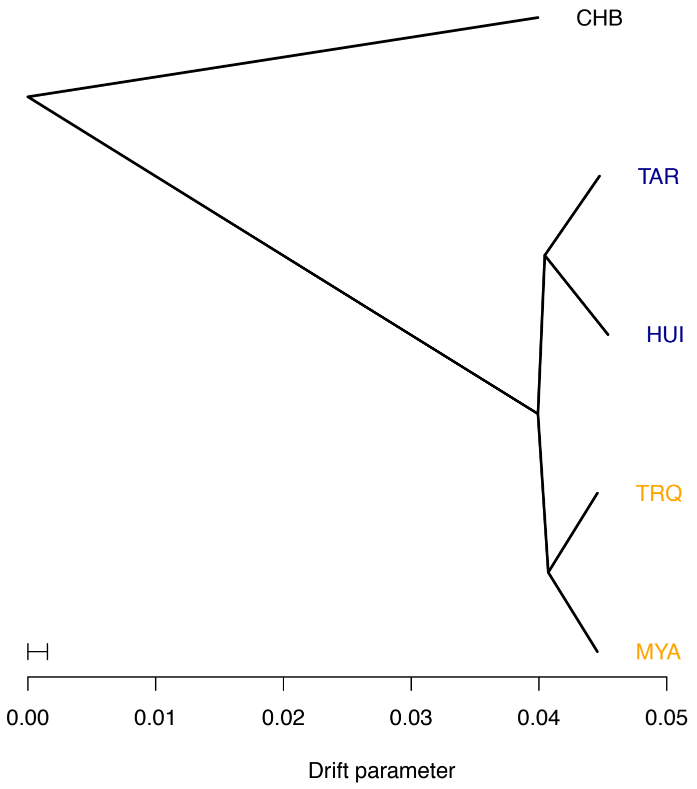


**Figure S6.**
Tree topology inferred by TreeMix. The tree groups together the two northern populations Rarámuri (TAR) and Huichol (HUI), and the two southern Triqui (TRQ) and Maya (MYA).

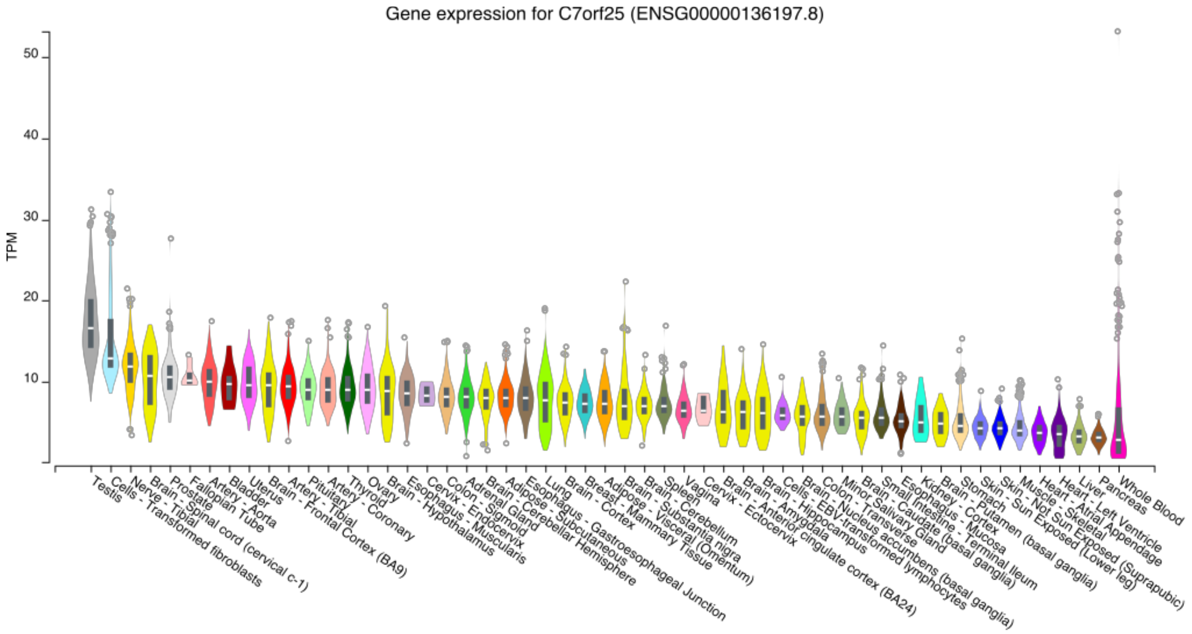


**Figure S7.**

Gene expression for *C7orf25*, a highly differentiated gene in Rarámuri. Tissues are shown in the x-axis, their expression is shown in the y-axis in Transcripts Per Million (TPM) units. Figure obtained from the GTEx Portal (GTEx Analysis Release V7) on 01/29/2019.

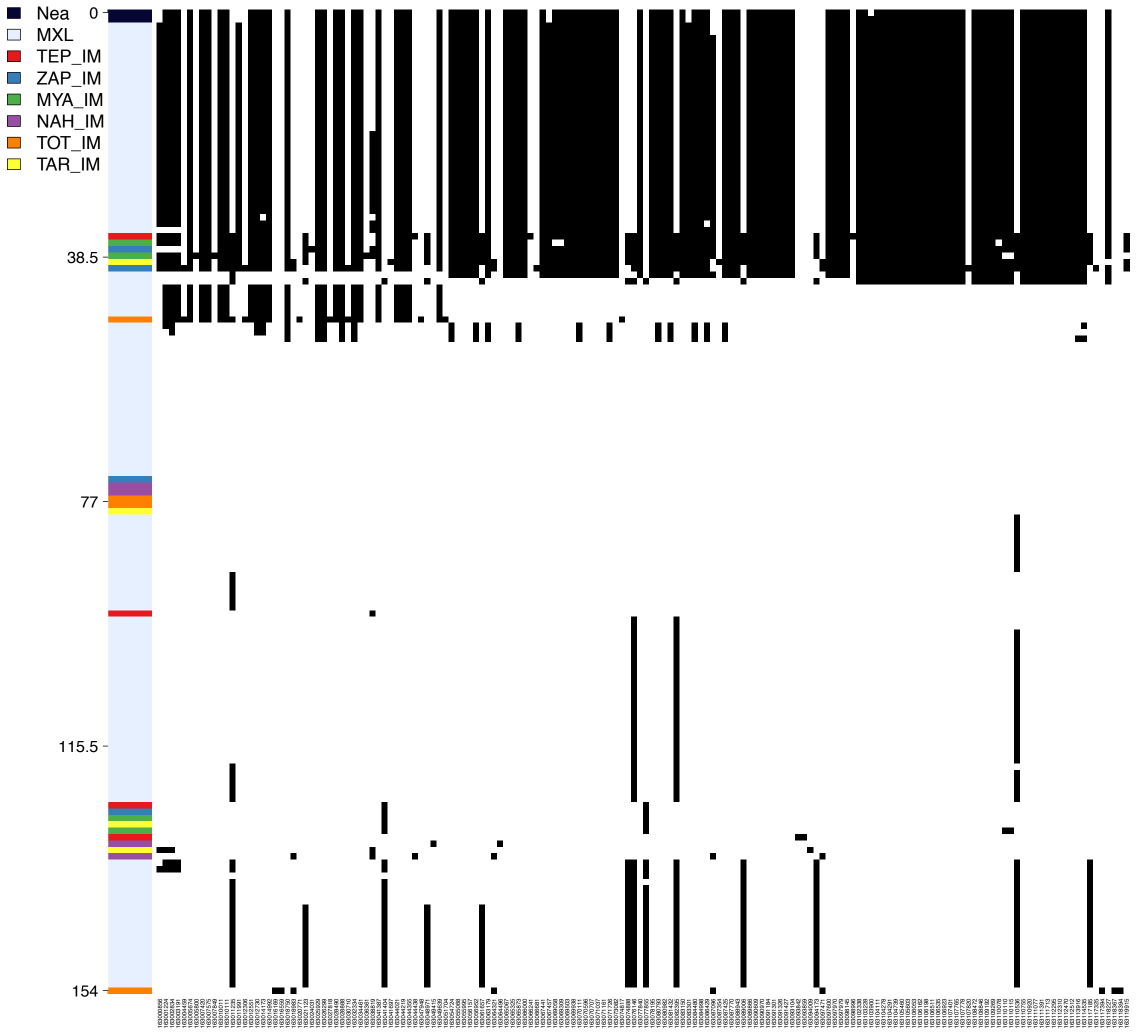


**Figure S8.**
Haplotype structure of *FAP/IFIH1* locus (chr2:163000000-163120000) obtained with the program Haplostrips (Marnetto and Huerta-Sánchez 2017) in Neandertal (Nea), Mexicans from the TGP (MXL), and NM. Derived alleles are shown in black. The top two haplotypes belong to Altai Neandertal. NM genomes in this figure were obtained from Romero-Hidalgo (2017). TEP: Tepehuano, ZAP: Zapotec, MYA: Maya, NAH: Nahua, TOT: Totonac, TAR: Tarahumara or Rarámuri.

#### Supplementary tables index

**Supplementary table S1.**
https://www.dropbox.com/s/t52lahq7aya6o3f/Table%20S1.xlsx?dl=0

**Supplementary table S2.**
https://www.dropbox.com/s/rdyccmtq1c1ixuf/Table%20S2.xlsx?dl=0

**Supplementary table S3.**
https://www.dropbox.com/s/081mvcbkw56t65i/Table%20S3.xlsx?dl=0

**Supplementary table S4.**
https://www.dropbox.com/s/3e8hhh6wwbjz7yg/Table%20S4.xlsx?dl=0

**Supplementary table S5.**
https://www.dropbox.com/s/uh6v79bo4n4f2am/Table%20S5.xlsx?dl=0

**Supplementary table S6.**
https://www.dropbox.com/s/io6bfes0j902u7v/Table%20S6.xlsx?dl=0

**Supplementary table S7-S10.**
https://www.dropbox.com/s/1m33s1fkh3r7juk/Table%20S7%20-S10.xlsx?dl=0

**Supplementary table S11.**
https://www.dropbox.com/s/lecpuk2jafosl3p/Table%20S11.xlsx?dl=0

**Supplementary table S12.**
https://www.dropbox.com/s/0klzmw46zoskull/Table%20S12.xlsx?dl=0

**Supplementary table S13.**
https://www.dropbox.com/s/cg0ewr0yyihrlxt/Table%20S13.xlsx?dl=0
